# Inhibition of NLRP3 by a CNS-penetrating indazole scaffold

**DOI:** 10.1101/2025.07.01.662566

**Authors:** Jane Torp, Dominic Ferber, Hannes Buthmann, Gregor Hagelueken, Aditi Deshpande, George Hartman, Robert E. Hughes, Taiz Salazar, Sarah Tronnes, Assem Duisembekova, Michael Marleaux, Inga V. Hochheiser, Rebecca C. Coll, Rusty Montgomery, Kristen Fortney, Bénédicte F. Py, Kevin Wilhelmsen, Matthias Geyer

**Affiliations:** Institute of Structural Biology, University of Bonn, Venusberg Campus 1, 53127 Bonn, Germany; BioAge Labs, 5885 Hollis St., Suite 370, Emeryville, CA 94608, USA; CIRI, Centre International de Recherche en Infectiologie, Univ Lyon, Inserm, U1111, Université Claude Bernard Lyon 1, CNRS, UMR5308, ENS de Lyon, Lyon, France; Wellcome-Wolfson Institute for Experimental Medicine, Queen’s University Belfast, Belfast BT9 7BL, UK

## Abstract

Low-grade inflammation is a hallmark of ageing and a key cause of age-related impairments and diseases^1^. The NOD-like receptor NLRP3 senses a variety of danger signals and environmental insults, resulting in pro-inflammatory response, inflammasome formation and pyroptosis^2,3^. Its aberrant activation has been linked to many acute and chronic diseases ranging from atherosclerosis to Alzheimer’s disease and cancer, making NLRP3 an attractive therapeutic target^4,5^. Here we report the discovery, characterization, and structure of an indazole-based NLRP3 antagonist, BAL-1516, which potently inhibits inflammasome formation in monocytes and microglia. The cryo-electron microscopy structure of BAL-1516 bound to NLRP3 reveals a previously undescribed compound binding site at a surface groove of the nucleotide-binding domain with contacts to the FISNA and WHD subdomains. The characteristic feature of BAL compound binding is the formation of three hydrogen bonds to the peripheral β-strand of the triple-ATPase; two from the indazole’s nitrogen atoms and a third from the compounds’ linker region. Additional phenyl and thiazole moieties render the compound hydrophobic, allowing excellent blood-brain barrier penetration. The compound binding site is highly specific for NOD-like receptors, and the optimized compound BAL-1516 is able to directly bind mouse NLRP3 despite two conservative residue changes in the binding interface. The BAL compounds represent a first-in-class family of NLRP3 inhibitors, providing a broad design space, including covalent and degradative properties, for the development of NLRP3-directed therapeutics. The innate immune system contains cytosolic proteins that sense cellular stress caused by bacterial, viral and fungal infections or sterile inflammation, to control cellular integrity^2^. NLRP3 is a well-studied member of the nucleotide-binding oligomerization domain (NOD)-like receptors (NLRs) that is involved in the activation of the inflammasome, a multiprotein complex that mediates inflammation^6^. Upon detection of stress or pathogen-associated signals, NLRP3 triggers the activation of caspase-1, which leads to the production of pro-inflammatory cytokines such as IL-1β and IL-18, driving inflammatory responses and ultimately pyroptotic cell death. In the context of neuroinflammation, NLRP3 plays a significant role in the pathogenesis of various neurodegenerative diseases, including Alzheimer’s disease, Parkinson’s disease, and multiple sclerosis^7^. Research into targeting NLRP3 signalling with CNS-penetrating molecules holds potential for developing therapeutic strategies to alleviate neuroinflammatory conditions and to slow the progression of neurodegenerative diseases.

---

Small molecule inhibitors targeting NLRP3 have been described including the sulfonylurea compound MCC950, also known as CRID3^8,9^. MCC950 was developed based on early observations from cell-based screens that identified diarylsulfonylureas, such as glyburide, as inhibitors of NLRP3 signalling^10^. Cryo-electron microscopy and X-ray crystallography structures revealed the binding specificity of this compound, targeting a deep crevice in between the regulatory NACHT domain and the sensory leucine-rich repeat (LRR) domain, with the sulfonylurea being sandwiched between two opposing arginines^11-13^. Inactive, human NLRP3 was shown to form a spherical ‘cage’ structure made of ten subunits, where the effector Pyrin domain (PYD) is shielded inside, preventing its interaction with the downstream ASC protein^11^. Activation of NLRP3 requires disassembly of the cage, possibly mediated by the NEK7 kinase, and a ∼90° rotation of the subdomains within the NACHT. This allows formation of a ‘disc’-like structure and exposure of the PYDs, which act as nucleation seed for ASC filament elongation^14,15^. The conformational change is catalysed by the nucleotide-binding domain (NBD), a member of the triple ATPase family, accompanied by an ADP-to-ATP exchange. Multiple autosomal mutations of the *NLRP3* gene have been associated with constitutively active protein variants, leading to chronic inflammation summarized as cryopyrin-associated periodic syndrome (CAPS), which destabilise the inactive conformation and are therefore less susceptible to MCC950 binding^16,17^.

Today, MCC950 derivatives optimized for stability and tolerability are in multiple clinical trials with indications ranging from CAPS syndromes to osteoarthritis, obesity, cardiovascular diseases and Parkinson’s disease^4,18^. Here we report the development and structure of a highly potent inhibitor based on an indazole scaffold that targets NLRP3 at a site adjacent to the central β-sheet of the triple ATPase but different from the sulfonylurea binding site.

## BAL-1516 binds to NLRP3 at a site different from MCC950

The lead compound^19^, BAL-0028, has been optimised through several rounds of modification to BAL-1516 to achieve higher affinity, specificity, and the ability to penetrate the blood-brain barrier. To accomplish this, a 4-methylthiazole replaces the 3-fluorophenyl subunit at the western side, while the eastern indazole moiety is modified to 5-azaindazole extended with a 4-methyl substituent (Fig. 1a). In addition, a chiral centre is introduced at the azaindazole benzylic site connecting it to the methyl-amino-carbonyl moiety. We characterized the interaction of BAL compounds with NLRP3 by thermal stability measurements using dynamic light scattering (DLS). For comparison, a derivative lacking the chiral centre, BAL-0898, and MCC950 were included in these measurements (Extended Data Fig. 1a). To 3 µM full length, wild-type NLRP3 protein in the ADP-bound decamer conformation, 10 µM compound was added and the hydrodynamic radius measured upon increasing temperature as an indicator of protein unfolding and aggregation (Fig. 1b and Extended Data Fig. 1b,c). The addition of MCC950 did not result in significant protein stabilization, whereas BAL-0898 and BAL-1516 stabilized NLRP3 unfolding by more than 2.5 degrees Celsius over the DMSO control, indicating that the BAL binding site is readily accessible even in the preformed, intact decamer (Extended Data Fig. 1d).

**Fig. 1.**
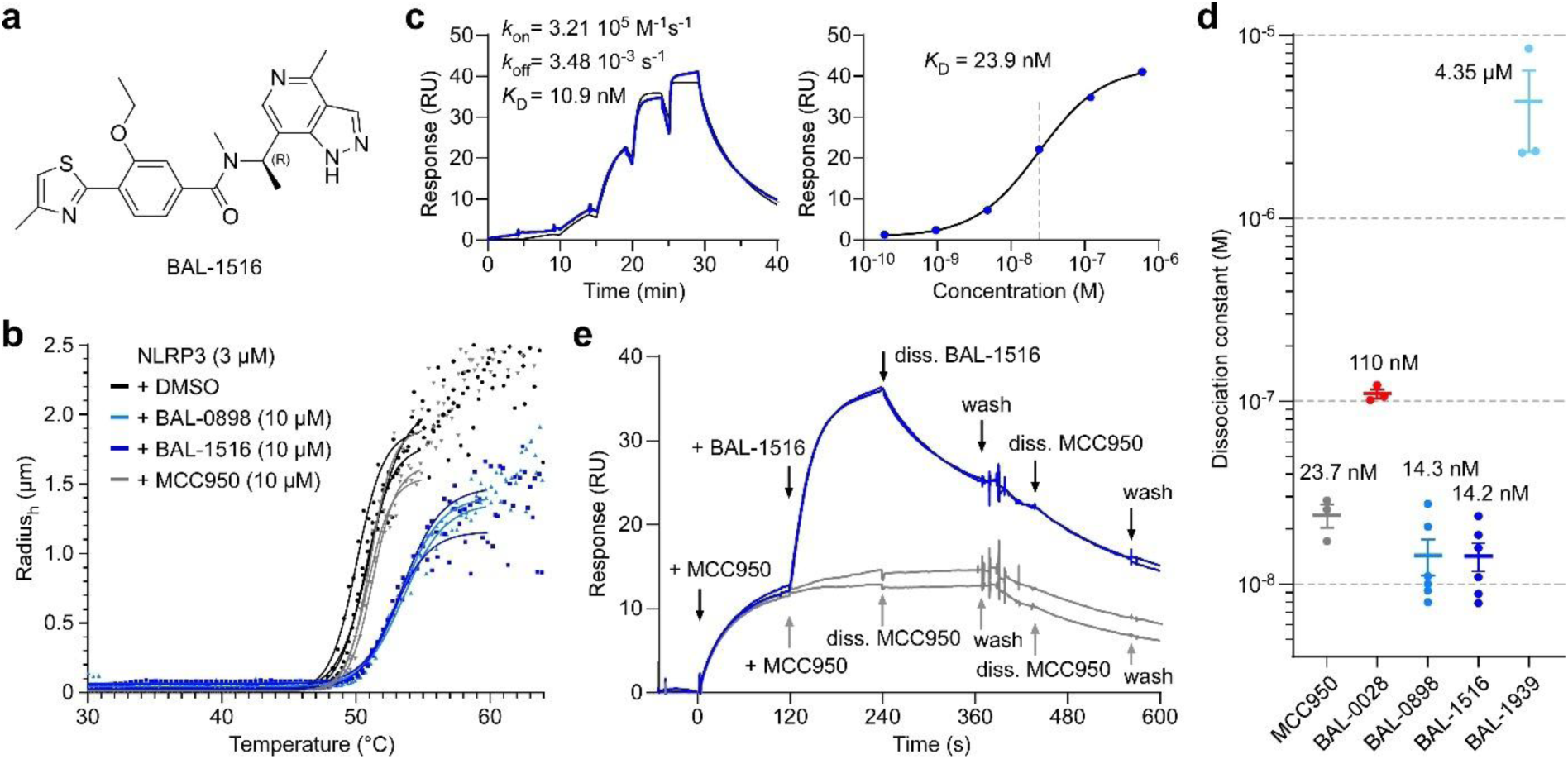
BAL-1516 binds with high affinity to NLRP3 at a site different from MCC950. **a**, BAL-1516 structure. **b**, Addition of BAL compounds 0898 or 1516 increases the thermal stability of NLRP3 over the MCC950 or the DMSO control. The hydrodynamic radius was determined by dynamic light scattering using 3 µM full length, wild type MBP-NLRP3 (peak 2) with 10 µM compounds upon temperature increase. **c**, Surface plasmon resonance of the interaction of BAL-1516 with NLRP3 reveals a dissociation constant (*K*_D_) of 10.9 nM. RU, response units. **d**, Equilibrium dissociation constants of MCC950, BAL-0028, BAL-0898, BAL-1516 and BAL-1939 as determined by SPR. The interaction of the enantiomer BAL-1939 is more than 300 times weaker than that of its eutomer BAL-1516, indicating the importance of this chiral centre. The average *K*_D_’s of *n* ≥ 3 independent experiments are given on top. **e**, BAL-1516 can bind on top of MCC950 to NLRP3, indicating its different binding site. SPR measurements were performed in A–B–A cycle mode, with MCC950 presented first (A), followed by BAL-1516 (B) and two successive dissociation steps (blue lines). Addition of MCC950 in cycle (B) is shown as control (grey lines).

Surface plasmon resonance (SPR) experiments were performed to determine the binding affinity of BAL compounds to NLRP3 (Fig. 1c,d and Extended Data Fig. 1e). Using the single cycle kinetics mode with immobilized protein, BAL-1516 showed a dissociation constant (*K*_D_) of 14.2 ± 2.5 nM to the NACHT domain of NLRP3. This indicates an 8-fold increase compared to the lead compound BAL-0028 (*K*_D_ = 110 ± 6 nM), and an affinity almost twice that of MCC950 (*K*_D_ = 23.7 ± 3.4 nM). In comparison, the (*S*)-enantiomer, BAL-1939, of the chiral centre connecting the phenyl ring with the indazole moiety revealed a *K*_D_ of 4.4 ± 2.1 µM; more than two orders of magnitude weaker binding than the eutomer. Using the A–B–A titration cycle with MCC950 (reservoir A) and BAL-1516 (reservoir B) we performed binning experiments of the compound binding site. The addition of BAL-1516 to pre-loaded MCC950 showed an increase in response units, indicating additive binding behaviour (Fig. 1e). In comparison, addition of MCC950 in the second cycle as a control showed no spectroscopic change. Reversing the compound titration scheme with BAL-1516 administered first followed by the addition of MCC950 resulted in a similar additive protein targeting behaviour (Extended Data Fig. 1f). The effect of BAL compounds on the intrinsic ATP-hydrolysis activity of NLRP3 was measured in HPLC multi-turnover assays^20^. Here, BAL-0898 and BAL-1516 did not affect nucleotide hydrolysis, indicating that the catalytic activity of NLRP3 remains unchanged by the compounds (Extended Data Fig. 1g). These data show that BAL-1516 binds with high affinity to NLRP3 at a site different from MCC950 without affecting ATP-hydrolysis activity, and in a specific configuration determined by the compound’s chiral centre.

## Structure of the NLRP3–BAL-1516 complex

Recombinant, full-length human NLRP3 (residues 3-1036) elutes in size-exclusion chromatography (SEC) experiments in two fractions: one of a high-molecular-weight assembly and one at the defined size of a decamer^11^. To investigate the effect of BAL-1516 on NLRP3 assembly, we added the compound either during the expression and gel filtration process or throughout the protein purification. Both procedures did not alter the elution profiles, indicating that the compound neither leads to aggregation nor disassembly of the protein complex (Extended Data Fig. 1b). Multiangle light scattering (MALS) combined with SEC at 280 and 325 nm wavelength confirmed the presence of the compound in the decamer assembly when adding the compound to the homogeneous sample (Extended Data Fig. 1c). A cryo-EM dataset of 6,106 micrographs was recorded on a Titan Krios microscope equipped with a Gatan K3 direct electron detector, imaging 1,739,268 particles. Multiple rounds of 2D classifications and refinements led to the selection of 765,958 particles for refinements (Methods). The NLRP3 decamer was reconstructed at an overall resolution of 3.7 Å in D5 symmetry (Extended Data Fig. 2). Focused refinement with a monomer mask in C1 symmetry led to a resolution of 3.06 Å for a single subunit (Extended Data Table 1). Initial model building was performed with chain A of the human NLRP3 complex structure^11^. A continuous amino acid chain was built from residues 133-1036, except for the acidic loop (686-725) and two gaps (178-199; 541-549) in the FISNA and HD2 subdomains. The N-terminal PYD effector domain was not resolved in the reconstructions.

The density of the BAL-1516 compound with a molecular mass of 435.6 Da was identified on the surface of NLRP3 in the nucleotide binding domain (Fig. 2a and Extended Data Fig. 3a). It is located at a site adjacent to the AAA+ ATPase defining β-sheet and opposite the bound ADP nucleotide (Fig. 2b,c and Extended Data Fig. 3b). The binding site covers a buried surface area of 620 Å^2^ of BAL-1516, which corresponds to 94% of its accessible surface area. Most direct interactions of NLRP3 to the compound are mediated by the NBD (19 out of 26 interacting residues according to PDBePISA^21^), with minor contributions from the FISNA and WHD subdomains (Fig. 2d). The binding site for MCC950 instead is empty, as expected (Extended Data Fig. 3c). Two cysteines are in the immediate vicinity to the BAL compound (Extended Data Fig. 3d). The first is C279, whose sulphate atom is in 3.5–3.6 Å distance to both a carbon of the ethoxy group and the methyl group of the central amino-carbonyl linker. The second is C514, which is at the tip of the hairpin loop of the antiparallel β-strands in the WHD and in about 13 Å distance to the methyl group of the chiral centre.

**Fig. 2.**
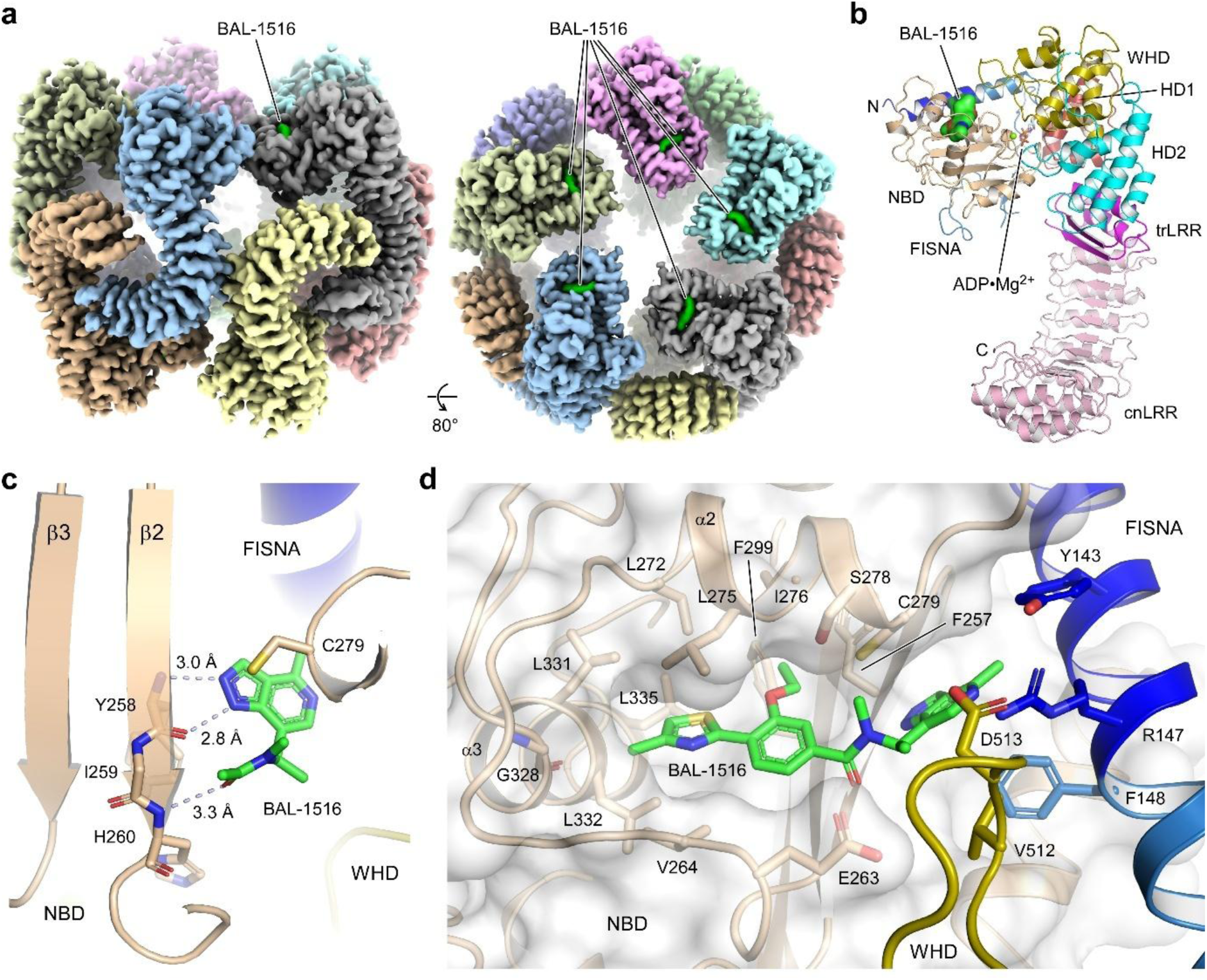
BAL-1516 binds to the triple-ATPase β-sheet in NLRP3. **a**, Structure of the NLRP3– BAL-1516 decamer in an equatorial (left) and apical (right) view. The cryo-EM density of a composite map is shown at a threshold level of 5.42 with the bound compound indicated (green density). **b**, BAL-1516 binds to the nucleotide-binding domain (NBD) of NLRP3. **c**, Binding of BAL-1516 is characterized by the formation of three hydrogen bonds between the 5-azaindazole ring and the aminocarbonyl linker of the compound and the main chain atoms of Y258 and H260 of strand β2 of NLRP3. Distances of the hydrogen bonds are indicated; the phenyl and thiazole rings are omitted for clarity. **d**, Molecular interactions of NLRP3 with BAL-1516. The binding site is mainly formed by a hydrophobic groove on the NBD and is supported by interactions with the FISNA and WHD subdomains.

## BAL-1516 interacts with the triple-ATPase β-sheet

The characteristic feature of BAL compound binding to NLRP3 is the formation of three hydrogen-bonds to the terminal β2 strand of the nucleotide-binding domain (Fig. 2c). Here, the 1*H*-indazole moiety interacts with the main chain atoms of Y258, while the carbonyl group in the compounds’ linker region forms a hydrogen bond with H260. This alternating acceptor– donor–acceptor pattern at ideal distances of 2.8–3.3 Å to the backbone atoms of NLRP3 is reminiscent of the formation of an antiparallel β-strand. The hydrophobic thiazole-phenyl ring system on the western side of the compound is framed by an extended loop between residues E263 and C279 of NLRP3 and contacts the central β-sheet of the NBD from behind the side of the ADP-Mg^2+^ nucleotide (Fig. 2d). The ethoxy group at the para-position of the phenyl ring sticks into a hydrophobic core formed by F257, I259 and F299 of the β-strands β2 and β3, and complemented by L272 and I276 from helix α2, which sits as a lid on top of the compound. The thiazole ring is rotated about 22° relative to the phenyl ring and is embedded by the side chains of residues V264, L331, L332, L335, L272, L275, F299 and I259, forming a hydrophobic cleft. The space to accommodate the 4-methyl group of the thiazole is provided by G328 at the beginning of helix α4. The chiral centre of BAL-1516 induces a kink between the indazole and the methyl-aminocarbonyl group, so that the compound with (*R*)-configuration fits precisely into the binding site on the surface of NLRP3. This observation could explain the about 300-fold higher affinity of BAL-1516 to NLRP3 compared to its (*S*)-enantiomer BAL-1939. The methyl group at the chiral centre points towards the surface of the protein-compound complex, providing an exit vector for molecule extensions (Extended Data Fig. 3d). The binding site appears rather uncharged in the surface display of the electrostatic potential (Fig. 3a), underlining the hydrophobic nature of the CNS-penetrating compound. In the decamer assembly of inactive NLRP3, the BAL-1516 interaction site is close to the apical axis in direct proximity to the basic cluster (131-147) of the FISNA, yet readily accessible from the bulk solvent (Extended Data Fig. 3e). A schematic of the interactions is shown in Fig. 3b.

**Fig. 3.**
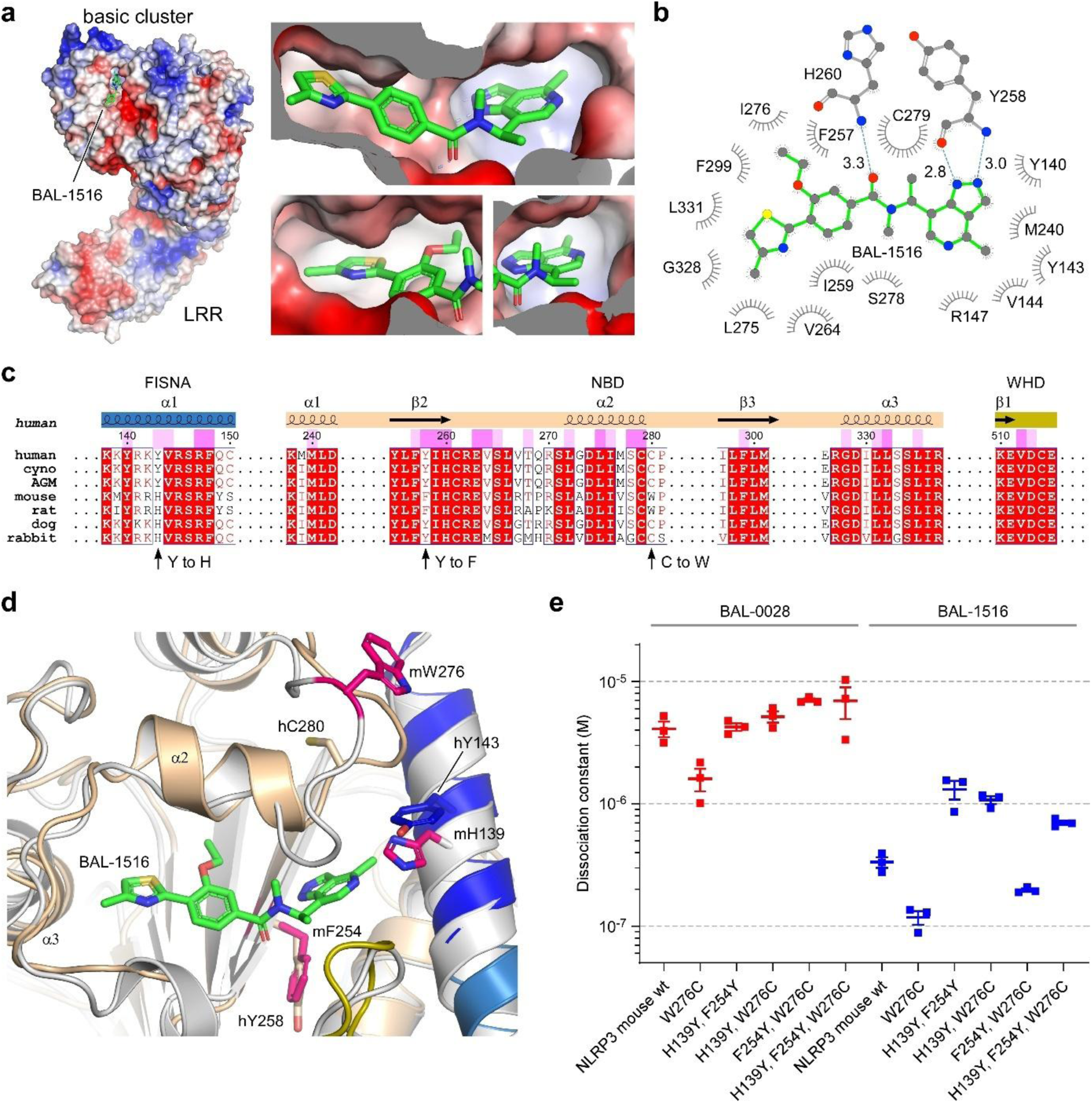
BAL-1516 binds mouse NLRP3. **a**, Electrostatic surface display of the BAL-1516 binding site in NLRP3 shows its proximity to the basic cluster while the thiazole, ethoxy and 5-azaindazole binding sites are largely hydrophobic. **b**, Lig-plot of the interactions between NLRP3 and BAL-1516. **c**, Sequence alignment of NLRP3 proteins from seven species tested for inhibition by BAL-0028. Residues interacting with BAL-1516 at 1–10 or > 10 Å^2^ buried surface area are marked with light or bright bars, respectively. **d**, Overlay of human (9IHN, this study) and mouse (7LFN)^31^ NLRP3 structures, displaying the amino acid differences in the BAL compound binding interface. **e**, SPR binding analyses of wild-type mouse NLRP3 or the proposed gain-of-function mutations with BAL-0028 and BAL-1516. None of the tested single, double and triple mutations significantly increased the *K*_D_ to BAL compounds, while the interaction with BAL-1516 was on average one magnitude better than binding to BAL-0028.

## BAL compound binding to mouse NLRP3 variants

Our initial studies with the lead compound BAL-0028 showed low efficacy for NLRP3 in mice, which severely limits its experimental potential for characterisations in aging and Alzheimer’s disease models^19^. Seven different species were tested, showing that mouse, rat, dog and rabbit NLRP3 had reduced susceptibility to BAL-0028 inhibition compared to cynomolgus, AGM and human. Knowing the binding site of the BAL compound, we identified two directly interacting residues that differ from their human counterparts; histidine at the position of Y143 and phenylalanine at the position of Y258 (Fig. 3c). A third alteration is the C280 change to tryptophan in mouse and rat NLRP3. Although at an indirect position, the significant change in size may impact the loops’ conformation and the preceding residue C279, which largely contributes to compound binding (Fig. 3d). We performed single, double and triple mutations of these three sites in mouse NLRP3 to generate gain-of-function mutations for BAL compound binding. SPR analysis showed that BAL-1516 binds to the wild-type mouse NLRP3 NACHT protein with a *K*_D_ of 200 nM, which is approximately one order of magnitude tighter than the binding of BAL-0028 (Fig. 3e). However, of the proposed gain-of-function mutants, only mouse W276C and the double mutant F254Y, W276C showed improved binding behaviour, whereas all mutant proteins containing the H139Y exchange appeared to be instable.

## BAL-1516 is a potent inflammasome inhibitor with CNS penetrance

The inhibitory potential of BAL-1516 on NLRP3-mediated inflammasomes was tested in HEK293T cells, monocytes, peripheral blood mononuclear cells (PBMCs) and microglia. First, we used HEK293T cells stably expressing BFP-ASC and doxycycline-inducible NLRP3 to monitor ASC specks by flow cytometry. Upon nigericin stimulation, a significant increase in ASC specks is observed, which is dose-dependently inhibited by BAL-1516, BAL-0898 and MCC950 with IC_50_ values of 14.5, 122 and 69 nM, respectively (Fig. 4a). We tested the synergy of BAL compounds and MCC950 in NLRP3 inhibition by comparing the composition of half the concentration of each compound with the use of either compound at full concentration. Three concentrations were analysed and BAL-0898 was used due to its better solubility. The reduction of ASC specks indicates similar apparent IC_50_ values, revealing an additive effect of both compounds with no significant influence on the binding affinity of the other antagonist (Fig. 4b). This observation agrees with the titration experiments by SPR and reflects the different binding sites of each compound class on the structure of NLRP3 (Fig. 4c), showing that neither over-additive effects due to increased binding affinity nor destructive effects due to decreased binding affinity of the other antagonist prevail.

**Fig. 4.**
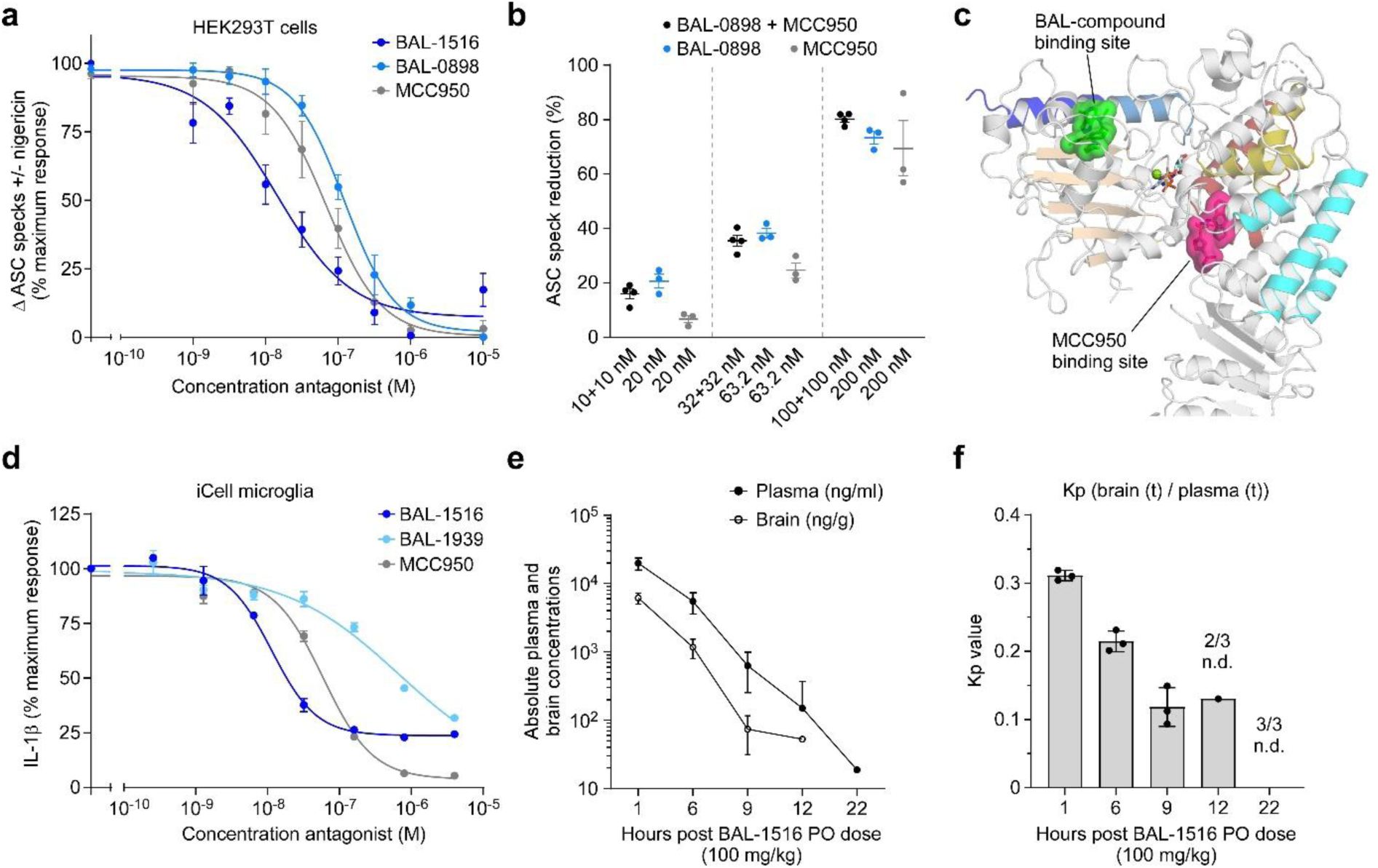
BAL compounds inhibit inflammasome activity in microglia. **a**, BAL-1516 is a potent inhibitor of ASC speck formation as assessed by flow cytometry in HEK293T ASC-BFP cells transfected with human NLRP3 and stimulated with nigericin. **b**, The inhibition of ASC speck formation in HEK293T cells by BAL-0898 and MCC950 at equal concentrations is additive. **c**, Display of the BAL compound and MCC950 binding sites in NLRP3. **d**, Dose response curves for BAL-1516, BAL-1939 and MCC950 inhibition of IL-1β release in iCell microglia. IC_50_ and IC_90_ values of 11 nM (55 nM), 673 nM (37.1 µM) and 59 nM (339 nM) for BAL-1516, BAL-1939 and MCC950, respectively, were determined from mean maximum response compared with positive control. Cells were primed with 200 ng/ml LPS for 4 h and stimulated with 10 µM nigericin for 30 min. **e**, Total plasma and brain concentrations of BAL-1516 were determined 1, 6, 9, 12 and 22 hr post oral dosing with BAL-1516 at 100 mg/kg in mice. Data plotted for mean ± s.e.m. of 2 independent experiments. **f**, The *K*_p_ value (the ratio of the total concentration of BAL-1516 in the brain to plasma) was determined for individual mice at 1, 6, 9, 12 and 22 hr post oral dosing with BAL-1516 at 100 mg/kg.

We compared BAL-0028 and BAL-0898 with MCC950 inhibition of NLRP3 activity in primary human monocytes and PBMCs. BAL-0028 and BAL-0898 show higher potency than MCC950 in inhibiting pyroptosis in LPS-primed monocytes treated with nigericin (Extended Data Fig. 4a), as well as IL-1β and IL-18 secretion in LPS-primed PBMCs treated with nigericin or monosodium urate crystals (MSU) (Extended Data Fig. 4b-e). We further evaluated BAL-1516 in induced pluripotent stem cell (iPSC)-derived microglia (iCell microglia) analysing the release of IL-1β and IL-18. BAL-1516 consistently inhibited LPS primed and nigericin stimulated iCell microglia in a dose-dependent manner for both cytokines at a higher potency than its BAL-1939 enantiomer (Fig. 4d and Extended Data Fig. 5a). IC_50_ values of 11 and 673 nM were determined for BAL-1516 and BAL-1939, respectively, for the reduction of IL-1β release compared to 59 nM for MCC950, and similar values were obtained for the inhibition of IL-18 release (Extended Data Fig. 5b). The cytotoxicity of BAL-1516 and BAL-1939 and their influence on cell viability was tested in iCell microglia in the same concentration range, showing no significant divergences in LDH release and cell functionality (Extended Data Fig. 5c,d). The ability for CNS penetration of BAL-1516 was analysed in mice by determining the absolute values of compound concentration in blood plasma and brain tissue over a time course of 22 hours after dose (Fig. 4e). The *K*_p_ coefficient as a measure for total brain concentration to total plasma concentration decreases from 0.3 to 0.1 from the first to the twelfth hour indicating excellent blood-brain barrier penetration and stability in tissue (Fig. 4f). These data confirm that BAL-1516 inhibits NLRP3 in microglia and exhibits CNS persistence.

## Discussion

Here we identify a yet unexplored targeting site on NLRP3 that allows for small molecular compound binding and highly potent inhibition of inflammasome formation. The binding site is located on the surface of the NACHT domain, primarily formed by the nucleotide-binding domain (NBD), with contributions from the FISNA and WHD subdomains. Moreover, the site is readily accessible in both, the spherical ‘cage’ structure and the concentrical ‘disc-like’ structure of the NLRP3 decamer. The compound targeting this site is built of an indazole linked to a hydrophobic biphenyl moiety and constitutes a chemical scaffold different from the sulfonylurea entity found in MCC950, in line with their different binding sites in NLRP3. Its hydrophobic nature allows excellent blood-brain barrier penetration, opening an avenue for the targeting of neuroinflammation related diseases.

Indazoles are rare in nature and considered a privileged scaffold in drug design^22^. The formation of three hydrogen-bonds, two from the indazoles’ nitrogen atoms and one following a chiral centre, is the key asset of BAL compound binding to NLRP3. In addition, the chiral centre contributes significantly to the compounds’ recognition specificity as the affinity of the corresponding enantiomer is more than 300-fold weaker. The chiral centre offers also an exit vector for the generation of degraders or fluorescently labelled compounds due to its solvent exposure and accessibility. The interaction to the peripheral β-strand of the triple-ATPase β-sheet delineates a unique binding site, also known as ‘Glu-switch’, as it provides a link between ATPase activity and ligand binding in AAA+ proteins^23^. In circular assemblies such as the chaperone p97 hexamer, the DNA clamp loader or the 26S proteasome, the Glu-switch mediates interaction to the neighbouring subunit, linking conformational changes to nucleotide turnover of the ATPase motor^24,25^. However, the formation of ‘disc-like’ structures for active NLRP3 or NLRC4 is mediated by the FISNA subdomain, rather than the β-sheet of the NBD^14,26^. While a similar binding site as for the BAL compounds in NLRP3 should be present in all NLRs, the primary sequence to its closest homologue, NLRP12, already differs in eight of 26 residues, resulting in no detectable interaction (Ext. Data Figs. 6 and 7). Recent advances to depart from the sulfonylurea class for better brain penetrability are NT-0796, an ester substituted carbamate derivative that acts as a pro-drug, or the pyridazine-based scaffolds NP3-253 and ASP0965, and a bicyclic variant thereof, which, however, all target the MCC950 binding site^27-30^.

The structure of the NLRP3–BAL-1516 complex provides initial insights into its mode of action as an inflammasome antagonist. The transition from inactive to active NLRP3 requires large conformational changes, including the decomposition of the LRR face-to-face interaction that constitutes the dimeric building block of the decamer assembly^11,14^. It is thought that the association to membranes is a requirement for this ‘opening-up’ of the structure and the compound binding site includes the basic cluster of the FISNA responsible for lipid binding^3^. The BAL compounds offer a new design space to explore NLR targeting and functional analyses. The determination of its binding site provides the structural framework for rational optimization and discovery of next-generation inflammasome therapeutics targeting NLRP3.

## Acknowledgments

The authors gratefully acknowledge the electron microscopy training, imaging and access time granted by the life science EM facility of the Ernst-Ruska Centre at Forschungszentrum Jülich (FZJ). We thank Thomas Heidler and Daniel Mann at FZJ for excellent help and advice on cryo-EM data acquisition and processing. We acknowledge access to the cryo-EM infrastructure of StruBiTEM (Cologne, funded by DFG Grant INST 216/949-1 FUGG) and excellent help from Monika Gunkel with plunge-freezing. We would like to thank the Flow Cytometry Core Facility of the Medical Faculty at the University of Bonn for providing help, services, and devices funded by the DFG–project number 216372401. We acknowledge the contribution of SFR Biosciences (Université Claude Bernard Lyon 1, CNRS UAR3444, Inserm US8, ENS de Lyon) and the help of the staff of LyMIC-PLATIM, as well as of the Etablissement Français du Sang Auvergne-Rhône-Alpes. This work was supported by a grant from the DFG to M.G. (GE 976/16-1). D.F. is supported by a fellowship from the Studienstiftung des deutschen Volkes, Bonn, Germany. A.Du. is supported by the Republic of Kazakhstan (Bolashak fellowship). B.F.P. is supported by the French National Research Agency (ANR-22-CE15-0032-01). M.G. is supported by the European Research Council (ERC Advanced Grant NalpACT) and by the DFG under Germany’s Excellence Strategy–EXC2151-390873048.

## Author contributions

J.T. performed protein expression and purification assays, cryo-EM recordings, and structure reconstructions with the help of Gr.H.; D.F. performed SPR binding experiments as well as DLS and MALS measurements; and H.B. conducted ATP hydrolysis measurements and ASC speck assays. A.De., T.S., S.T., A.Du. and B.F.P. performed cell-based potency assays; Ge.H., K.W. and R.E.H. contributed to the development of BAL-0028, BAL-0898 and BAL-1516; A.De. and K.W. designed and supervised research; M.G. designed and supervised research and wrote the manuscript. All authors discussed and commented on the manuscript.

## Methods

### Protein expression and purification

For biochemical and structural analyses human full length NLRP3 (wt; aa 3-1036; UniProt accession code Q96P20) was expressed with an N-terminal MBP affinity tag in *Sf9* insect cells. Human NLRP3-NACHT (wt; aa 131-694) and mouse NLRP3-NACHT (wt; aa 127-692; UniProt Q8R4B8), mouse NLRP3-NACHT (H139Y; aa 127-692), mouse NLRP3-NACHT (F254Y; aa 127-692), mouse NLRP3-NACHT (W276C; aa 127-692), double and triple mutants thereof, and human NLRP12-NACHT protein (wt; aa 122-679; UniProt P59046) were expressed with N-terminal Avi and MBP affinity tags in *Sf9* insect cells for SPR measurements. 1 L expression culture was infected with the respective 3% baculovirus preparation and incubated for 60 h after which the cells were harvested. The cell pellet was snap frozen in liquid nitrogen and stored at -80 °C until further use.

### Purification of the NLRP3-NACHT proteins for SPR

The cell pellet was resuspended with lysis buffer (50 mM HEPES pH 7.5, 150 mM NaCl, 10 mM MgCl_2_, 1 mM ADP, 5 mM β-mercaptoethanol) and supplemented with 1 mM PMSF and 1 µg/ml DNase I. The resuspended cells were lysed by sonication with 40% intensity, 4 min of 5 s on / 5 s off cycles, followed by centrifugation at 75,600 x *g* at 10 °C for 1 h. The supernatant was filtered through a 0.8 µm filter and applied to a MBPtrap (Cytiva) column, pre-equilibrated with lysis buffer and coupled to an ÄKTA Start system. Unbound proteins were washed off by 10 column volumes (CV) lysis buffer, followed by an elution of bound protein with lysis buffer supplemented with 15 mM maltose and 150 mM L-arginine. The fractions containing protein were pooled and concentrated with a 50 kDa MWCO Amicon^®^ Ultra (Merck) filter. The protein was subjected to a HiLoad 16/600 Superdex 200 pg column (Cytiva) equilibrated with SEC buffer (50 mM HEPES pH 7.5, 150 mM NaCl, 10 mM MgCl_2_, 1 mM ADP, 0.5 mM TCEP, 150 mM L-Arginine). Protein eluting between 70-80 ml for human NLRP3 or between 65-80 ml for mouse NLRP3 proteins were pooled and concentrated with a 50 kDa MWCO Amicon^®^ Ultra (Merck) filter to a concentration of 1.1-2.0 mg/ml, snap frozen in liquid nitrogen and stored in aliquots at -80 °C until further use.

### Purification of the human full-length NLRP3 protein

The cell pellet was resuspended with lysis buffer (50 mM HEPES pH 7.5, 150 mM NaCl, 10 mM MgCl_2_, 1 mM ADP, 0.5 mM TCEP) containing 10 µM BAL-1516 compound and supplemented with 1 mM PMSF. The resuspended cells were lysed by sonication with 40% intensity, 4 min of 5 s on /5 s off cycles, followed by centrifugation at 75,600 x *g* at 10 °C for 1 h. The supernatant was filtered through a 0.8 µm filter and applied to a MBPtrap (Cytiva) column, pre-equilibrated with lysis buffer and coupled to an ÄKTA Start system. Unbound proteins were washed off by 10 CV lysis buffer, followed by an elution of bound protein with lysis buffer supplemented with 15 mM maltose.

For cryo-EM analysis, fractions containing protein were pooled and concentrated with a 100 kDa MWCO Amicon^®^ Ultra (Merck) filter to 2.32 mg/ml. The MBP affinity tag was cleaved off by adding 1:50 ratio of in-house produced TEV protease and incubated at 4 °C o/n. Prior to size exclusion chromatography, the protein was cross-linked with BS3 cross-linker (ThermoFisher). For this, the TEV-cleaved protein was centrifuged at 10,000 x *g* and 4 °C for 5 min. The supernatant was incubated with 0.5 mM BS3 at 30 °C and 300 rpm for 30 min, followed by a quenching reaction with 95 mM (NH_4_)HCO_3_ for 15 min at 30 °C and 300 rpm. The cross-linked protein was centrifuged at 10,000 x *g* and 4 °C for 10 min, and 500 µl was applied to a Superose 6 increase 10/300 GL column (Cytiva), pre-equilibrated with lysis buffer. The fractions eluting between 12.5-14.0 ml (peak 2) were pooled and concentrated with a 100 kDa MWCO Amicon^®^ Ultra-0.5 (Merck) to 0.78 mg/ml and used for plunge freezing.

For ATPase activity measurements, the cell pellet from 2 l cell expression of MBP-tagged NLRP3 (f.l., wt) was resuspended and purified as described above and concentrated to 6.02 mg/ml. Protein was centrifuged at 10,000 x *g* for 10 min at 4 °C and 2 ml was applied to a Superose 6 prep grade XK 16/70 column (Cytiva), pre-equilibrated with lysis buffer in the absence of ADP. Fractions eluting between 41.0-48.0 ml (peak 1) were pooled, concentrated with a 100 kDa MWCO Amicon Ultra (Merck) filter to 0.97 mg/ml, snap frozen in liquid nitrogen and stored at -80 °C until further use.

### Cryo-EM grid preparation and data acquisition

Freshly purified protein was plunge frozen at the StruBiTEM facility of the University of Cologne using a VitroBot MarkIV (ThermoFischer) plunge freezer operating at 4 °C, 100% humidity with 3 s blot time. 3 µl of protein sample was applied to a plasma treated R1.2/1.3 Cu 300 mesh Holey carbon grid (Quantifoil) and plunge frozen into liquid ethane at liquid nitrogen temperature.

Cryo-EM micrographs were recorded with a Titan Krios microscope running at 300 kV and equipped with Gatan BioContinuum imaging filter and Gatan K3 camera (energy slit width 20 eV) at the cryo-EM facility of the Research Centre Jülich. 6,106 micrographs were collected at 105,000x nominal magnification, corresponding to a pixel size of 0.82 Å. The defocus range was -2.2 to -0.8 µm and electron exposure rate was 17.9 e^-^/Å^2^/s with an exposure time of 2.78s. The total electron exposure was 51.7 e^-^/Å^2^ over 65 frames.

### Cryo-EM data analysis

Micrographs were gain- and motion-corrected with Warp^32^. All further data analysis was done using CryoSPARC v4.4.0. Patch CTF job was used to estimate the contrast transfer function. Particle picking and multiple rounds of 2D classifications yielded 858,377 particles, which were used for ab-initio reconstruction with 2 classes followed by heterogeneous refinement. Next, the best class containing 765,958 particles was further refined with homogeneous and non-uniform refinement jobs yielding an overall GSFSC resolution of 3.66 Å at 0.143 FSC. Finally, local refinement was done with a mask on the best-resolved monomer, yielding GSFSC resolution of 3.06 Å at 0.143 FSC. For visualization purposes, the map from the local refinement was improved with DeepEMhancer (highRes)^33^.

### Model building

One monomer of human NLRP3 (PDB ID: 7pzc) was used as a starting model for building the protein chain. The amino acid positions were fitted to the electron density map with ChimeraX v1.6 and ISOLDE^34^ using the map from the local refinement. The final refinement was performed with PHENIX real-space refinement^35^.

For the decamer map and model, a composite map was generated from the local refinement map using Phenix Combine Focused Maps. For this, D5 symmetry was applied in the non-uniform refinement job and used as a target map, to which the locally refined monomer maps and models were aligned. The visual representations of the electron density map were generated using ChimeraX and PyMol. The cryo-EM maps and the respective models are deposited in the Electron Microscopy Data Bank (EMDB) and the Protein Data Bank (PDB), respectively.

### Surface plasmon resonance spectroscopy

SPR experiments were performed using a Biacore 8K instrument (GE Healthcare) operated at 25 °C and flushed with running buffer (10 mM HEPES pH 7.4, 200 mM NaCl, 0.5 mM ADP, 0.5 mM tris(2-carboxyethyl) phosphine (TCEP), 2 mM MgCl_2_, 1 g/l carboxymethyl dextran hydrogel (CMD), 0.05 % Tween20, 2 % DMSO). Avi-MBP-tagged human NLRP3^NACHT^ protein (131–694) was coexpressed with BirA in *Sf9* insect cells to achieve biotinylation. After purification by affinity and size-exclusion chromatography the biotinylated protein was immobilized on a streptavidin sensor chip (Cytiva). Following equilibration (100 µl min^-1^) for at least 2 h, different concentrations of test compounds were injected in single-cycle kinetics mode (30 μl min^-1^, association phase 240 s, dissociation phase 360 s) or using a customized A-B-A cycle method (30 µl min^-1^, 120 s injection of analyte A, 120 s injection of analyte B, 120 s injection of analyte A, wash, 120 s wait). Data were collected at a rate of 10 Hz. The binding data were double referenced by reference flow cell subtraction and blank cycle. Data were corrected by a 4-point DMSO solvent correction. To study BAL compound binding to mouse NLRP3 an Avi-MBP-tagged NLRP3^NACHT^ protein (127–692) or the respective mutant protein was immobilized. For NLRP12 SPR analysis the human Avi-MBP-tagged NLRP12^NACHT^ protein (122–679) was used. Processed data were fitted to a 1:1 interaction model using the Biacore Insight Evaluation Software (version 6.0.7.1750, Cytiva).

### DLS measurements

All samples were prepared with 3 µM MBP-tagged NLRP3 (f.l., wt) peak 2 centrifuged at 10,000 x *g* for 10 min and 10 µM BAL compound (final DMSO concentration: 0.1%). The negative control contained 0.1% DMSO instead of a compound. Using single use cuvettes (Wyatt Technology) the samples were measured in a DynaPro NanoStar DLS (dynamic light scattering) instrument (Wyatt Technology) at a heating rate of 0.5 °C/min starting at a temperature of 25 °C (final temperature: 75 °C). For every compound, at least two independent measurements of 3 single data acquisitions per temperature with acquisition times of t = 5 s were recorded. Data was evaluated using the DYNAMICS software (version 7.8.3.15, Wyatt Technology).

### ATP-hydrolysis assay

For the HPLC-based ATP-hydrolysis assay, 3 µM MBP-NLRP3 (f.l., wt) peak 1 was incubated for 30 min on ice in presence of 2% DMSO or 10 µM BAL compound. After incubation, 100 µM of ATP was added and the reaction was incubated for 68 min at 25 °C. Every 10 min a 10 µl sample was injected onto a 1260 Infinity II LC system connected to a reversed phase C18-silica column (Chromolith Performance, Merck) that was pre-equilibrated with buffer (30 mM K_2_HPO_4_, 70 mM KH_2_PO_4_, 10 mM TBA-Br, 4% v/v acetonitrile; pH 6.5). The detector was setup to measure at 259 nm wavelength. Detected peaks for ADP and ATP were integrated to calculate the molar concentrations of educt and product.

### SEC-MALS measurements

Size-exclusion chromatography coupled to multiangle light scattering (SEC-MALS) measurements were performed on an Infinity liquid chromatographic system (Agilent Technologies) equipped with a 1260 Infinity G5611A Bio-inert Quaternary Pump (with inline degasser), a 1260 Infinity G5667A Bio-inert High Performance Autosampler, a 1290 Infinity G1330B Thermostat and a 1260 Infinity G1365D Multiple Wavelength Detector coupled to a miniDAWN MALS device and an Optilab T-rEX RI detector (both Wyatt Technology). For the SEC-MALS measurements an analytical Superose 6 5/150 column (Cytiva) was used. The flow rate was set to 0.5 ml min^-1^. The chromatograms were recorded under isocratic conditions using 100% NLRP3-SEC-MALS buffer (50 mM HEPES pH 7.5, 200 mM NaCl, 10 mM MgCl_2_, 1 mM ADP, 0.5 mM TCEP, 1 g/l CMD). Each sample contained 10 μM of the respective protein and the indicated concentration of the respective compound (stock solutions in DMSO). The total volume of each sample was 60 μl, the concentration of DMSO in all samples was 2%. The negative controls contained 2% DMSO instead of a compound.

### Sequence alignment and binding site analysis

Sequence alignment of NLRP3 proteins from human (UniProt accession number Q96P20), mouse (Q8R4B8), rat (D4A523), rabbit (G1SNY9), cynomolgus (*macaca fascicularis*; A0A2K5WM90), African green monkey (AGM, *chlorocebus sabaeus*; A0A0D9R9G3), and dog (A0A8I3P020) species was performed with MultAlin, and the degree of conservation and the secondary structure annotated with ESPript^36^. The binding interface of BAL-1516 with human NLRP3 was determined with PDBePISA^21^ and marked with bars above the sequence.

### ASC speck formation assay

HEK293T cells stably expressing ASC-BFP fusion protein were used as previously described^11^. Cells were seeded into 24-well TC plates at a density of 125,000 cells per well and incubated overnight at 37 °C. To induce moderate expression of NLRP3 for assessable ASC speck formation, cells were transfected with 100 ng per well of a doxycycline inducible TetO6-NLRP3-hPGK-TetON3G-T2A-mCherry construct using Lipofectamine2000 (ThermoFisher Scientific) according to the manufacturer’s instructions. Sixteen hours post transfection, NLRP3 expression was induced by adding doxycycline (10 ng/ml). Simultaneously, cells were treated with increasing concentrations of BAL-1516, BAL-0898 or MCC950 and incubated for 4 h. To induce ASC speck formation, cells were subsequently treated with 10 μM nigericin for 1 h.

After trypsinization cells were washed and resuspended in flow buffer (DPBS, 2 mM EDTA, 0.5% BSA). Flow cytometry was conducted at a LSRFortessa II device (BD Biosciences). Gates were set to select for single cells expressing ASC-BFP. From these, mCherry-positive, NLRP3 expressing cells were selected. Among the mCherry-positive cells, the proportion of ASC speck formation was calculated as previously described by Sester et al. (2015)^37^. The difference in ASC speck formation between nigericin-stimulated and unstimulated cells was plotted against increasing antagonist concentrations. These values were normalized to the speck formation difference calculated for cells without compound treatment (% maximum response). The generated dose-response curve was fitted using the four-parameter logistic equation built into Prism 10.4.1 to determine the half maximal inhibitory concentrations of BAL-1516, BAL-0898 and MCC950.

### Human monocyte pyroptosis assay

Monocytes were purified from 10 ml of blood samples (drawn the day before in heparin tubes) by negative selection using the EasySep Direct Human Monocytes Isolation kit (StemCell) and plated at 10^5^ cells/ml in RPMI-1640 GlutaMax-1 without phenol red supplemented with 1x PS and 10% FBS (Gibco). Cells were primed with 40 ng/ml LPS (ultrapure, Invivogen) for 4 h and then treated with BAL compounds, MCC950, or vehicle only (DMSO, final concentration 0.05 %) for 15 min before addition of 5 µg/ml nigericin (Invivogen) and time-lapse imaging using CQ1 high content screening microscope (Yokogawa). PI (1.25 μg/ml) and Hoechst (0.2 μg/ml, both from Immunochemistry Technology) were added 2 h before imaging. Two images/well were taken every 15 min for 2 h using 10Å∼ objectives (UPLSAPO 10X/0.4). Images were quantified using the CQ1 software (Yokogawa) as described^16^. Briefly, the analysis was based on the “Total count in individual object” analysis module, which consists of reducing noise (Mean-Image, mask size 1 μm), thresholding the image (threshold gray level 150), removing pixels from the edges (Opening-Circle, 5 μm), separating adjacent nuclei (Find-Maximum-Distance with Minimum-Point-Distance 4 μm and Remove-Size 0.1 μm, then Dilation-Circle 3 μm, and Divide-Each-Region), integrating these last results with the intensity thresholding results (Expand-Region-3D) and applying a final filter size (50–500 μm). Pyroptosis was quantified by calculating the area under curve from the PI incorporation (%) over time curve of each well using Prism10.

### Human PBMCs IL-1β and IL-18 assay

PBMCs were purified from 10 ml of blood samples (drawn the day before in heparin tubes) by centrifugation on lymphocytes separation medium (Eurobio) at 800g for 20 min without brake, and plated at 10^6^ cells/ml in RPMI-1640 GlutaMax-1 supplemented with 1X PS and 10% FBS (Gibco). Cells were primed with 10 ng/ml LPS for 3 h, and then treated with BAL compounds, MCC950, or vehicle only (DMSO, final concentration 0.05 %) for 15 min before addition of 5 µg/ml nigericin for 2 h or 100 µg/ml MSU (Invivogen) for 3 h. Cell-free media were analysed using human IL-1β Duoset ELISA kit (RnD Systems) and ELISA using anti-IL-18 (D044-3, Biotechne) and anti-IL-18-biotin (D045-6, Biotechne) as capture and detection antibodies.

### Human iCell Microglial NLRP3 inflammasome activation assay

Human induced-pluripotent stem cell-derived microglia (iCell Microglia; R1131, Fujifilm) were directly thawed in poly-L-lysine coated 96-well plates at 50,000 cells/well in iCell glial base medium supplemented with iCell microglia supplements A, B, and C as per manufacturer’s instructions. Cells were grown in a humidified incubator at 37 °C, 5% CO_2_. Next day, 50% of the glial medium was replaced with fresh medium and cells were incubated for a further 2 days. For potency determination, iCell Microglia were first primed with 200 ng/ml LPS for 4 h and then pre-incubated for 30 min with 5-fold compound dilutions ranging from 0.256 nM to 4 µM. Cells were then stimulated with 10 µM nigericin with the corresponding compound concentrations and incubated for an additional 30 min. Supernatants were collected for cytokine ELISAs and to determine the level of cytotoxicity using the CytoTox 96® Non-Radioactive Cytotoxicity Assay kit (Promega, Madison, WI) according to the manufacturer’s supplied instructions. The remaining cells were processed for relative viability using the CellTiter-Glo® 3D Cell Viability Assay Kit (Promega, Madison, WI) according to the manufacturer’s supplied instructions. Cells treated with 200 ng/ml LPS and 10 µM nigericin (positive control) and cells with medium alone (negative control) were included in the experiment. MCC950 was used as a tool compound to compare relative potencies. For cytotoxicity (LDH release) calculations, raw absorbance values at 490 nm (OD_490_) of cell culture supernatants were normalized relative to the OD_490_ of the maximal LDH release control (after the addition of 10X Lysis Reagent for 30 minutes). Percent cytotoxicity was calculated using the formula, %Cytotoxicity = 100 X (experimental LDH release (OD_490_) / maximal LDH release (OD_490_)).

### In vivo BAL-1516 plasma and brain tissue concentration assays

Male CD-1 mice were orally dosed at 100 mg/kg with a 10 mg/ml solution of BAL-1516 in 20% hydroxypropopyl-β-cyclodextrin in water with a final pH of 3.96. At 1-, 6-, 9-, 12- and 22-hour post dose, plasma in K2-EDTA was collected from groups of three mice and the animals perfused prior to brain collection and homogenization with methanol: 15 mM PBS (1:2). BAL-1516 plasma and brain homogenate concentrations were determined using LC-MS/MS. The lower limit of quantification (LLOQ) was 1.00 ng/ml for plasma and 10.0 ng/g for brain. The Kp value was calculated for individual mice and is the ratio of total brain BAL-1516 (ng/g) to total plasma BAL-1516 (ng/ml).

### Study approval

Anonymous healthy donors’ blood was provided by the Etablissement Français du Sang (EFS) in the framework of convention #14-1820 between Inserm and EFS. Informed consent was received from participants prior to inclusion in the study.

### Data availability

The cryo-EM density reconstructions and models were deposited with the Electron Microscopy Data Bank (EMDB) (accession codes EMD-52874 for the NLRP3–BAL-1516 monomer and EMD-52874 for the NLRP3–BAL-1516 decamer) and with the Protein Data Bank (PDB) under accession code 9IHN (monomer) and 9Q8V (decamer). All data are available in the article or its supplementary files. Source data are provided with this paper.

---

**Extended Data Fig. 1.**
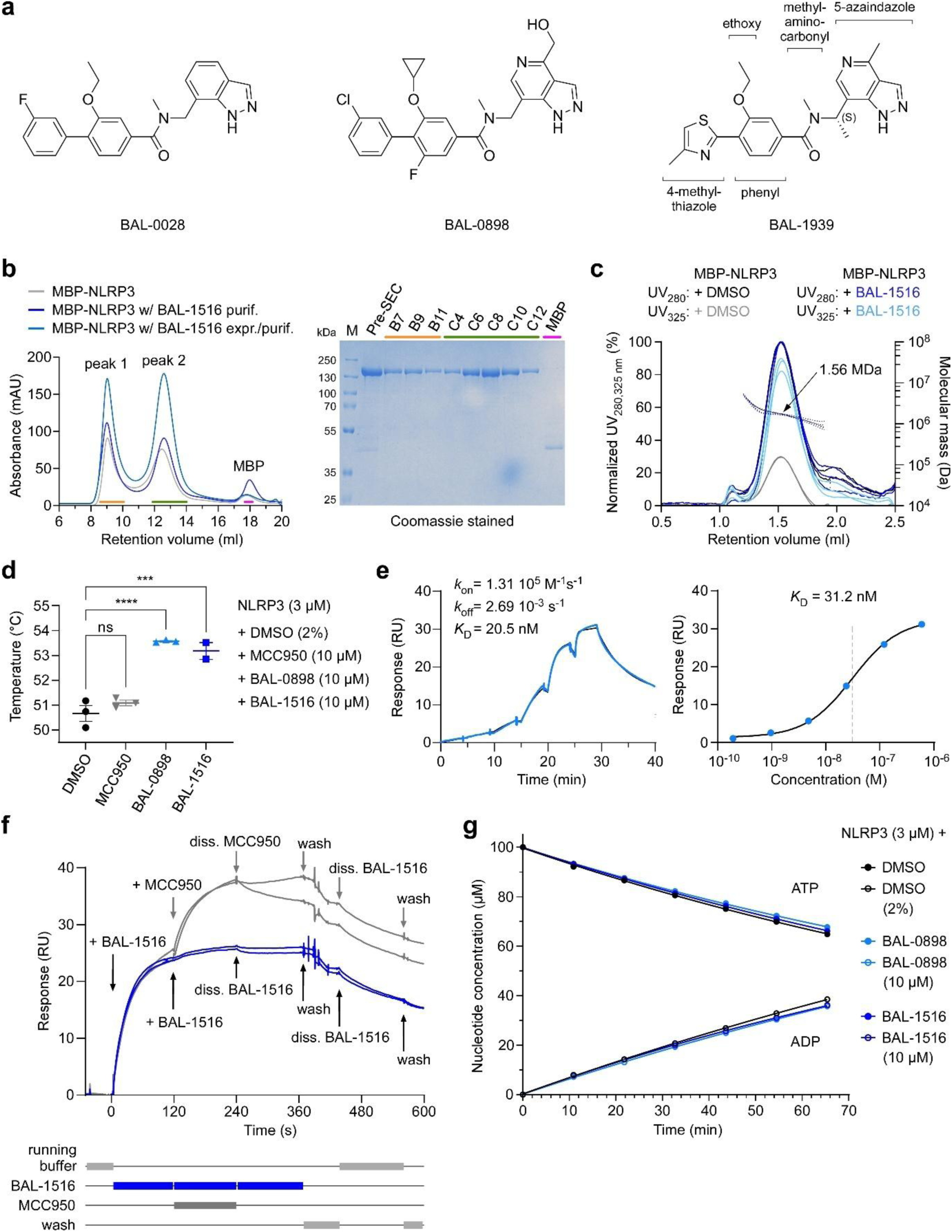
Biochemical characterization of BAL-1516 binding to human NLRP3. **a**, Structures of compounds BAL-0028, BAL-0898 and BAL-1939 used in this study. **b**, Size exclusion chromatography (left) and SDS-PAGE analysis (right) of human MBP-NLRP3 (f.l., wt) purified and/or expressed in the presence or absence of BAL-1516. The separation of NLRP3 into two peaks on a Superose 6 increase 10/300 column remains unchanged without (grey line) or with (blue line) 10 µM BAL-1516. The SDS PAGE shows MBP-NLRP3 with BAL-1516 present during the purification procedure. **c**, SEC-MALS experiment of 10 µM MBP-NLRP3 peak 2 with or without 30 µM BAL-1516 at a chromatography wavelength of 280 and 325 nm. The increase of the UV absorbance at 325 nm from the DMSO control (grey) to the sample with BAL-1516 (light blue) indicates that the compound migrates with the protein. **d**, Melting temperatures of NLRP3 upon addition of MCC950, BAL-0898 or BAL-1516 as determined by DLS for MBP-tagged (f.l., wt) peak 2 protein. Data are mean ± s.e.m. of n = 3 independent experiments (ns (not significant) *P* > 0.05, *** *P* < 0.001, **** *P* < 0.0001) (two-way ANOVA with Tukey’s multiple comparisons test). **e**, SPR measurement of the interaction of BAL-0898 with NLRP3 reveals a dissociation constant (*K*D) of 31.2 ± 4 nM. RU, response units. **f**, MCC950 binds in addition to BAL-1516 to NLRP3. SPR measurements of NLRP3 (131-694) with BAL-1516 and MCC950 in the A–B–A titration cycle. The additional association kinetics of MCC950 (B) after 120 s to the preformed NLRP3–BAL-1516 complex (A) indicates different binding sites for both compounds. **g**, BAL compounds 0898 and 1516 do not affect the intrinsic ATP-hydrolysis activity of NLRP3. Time-course experiment of a multi-turnover ATP-hydrolysis assay using MBP-NLRP3 (wt, f.l.) peak 1 at 3 μM concentration with either 10 µM BAL-0898 or 10 µM BAL-1516 or DMSO control incubated at 25 °C for 68 min in the presence of 10 mM MgCl2 and 100 μM ATP. The amount of ATP and ADP was determined in 10 min intervals using reverse phase HPLC. The eluted peaks were evaluated by integration. Shown is the mean ± s.e.m. of 3 independent experiments.

**Extended Data Fig. 2.**
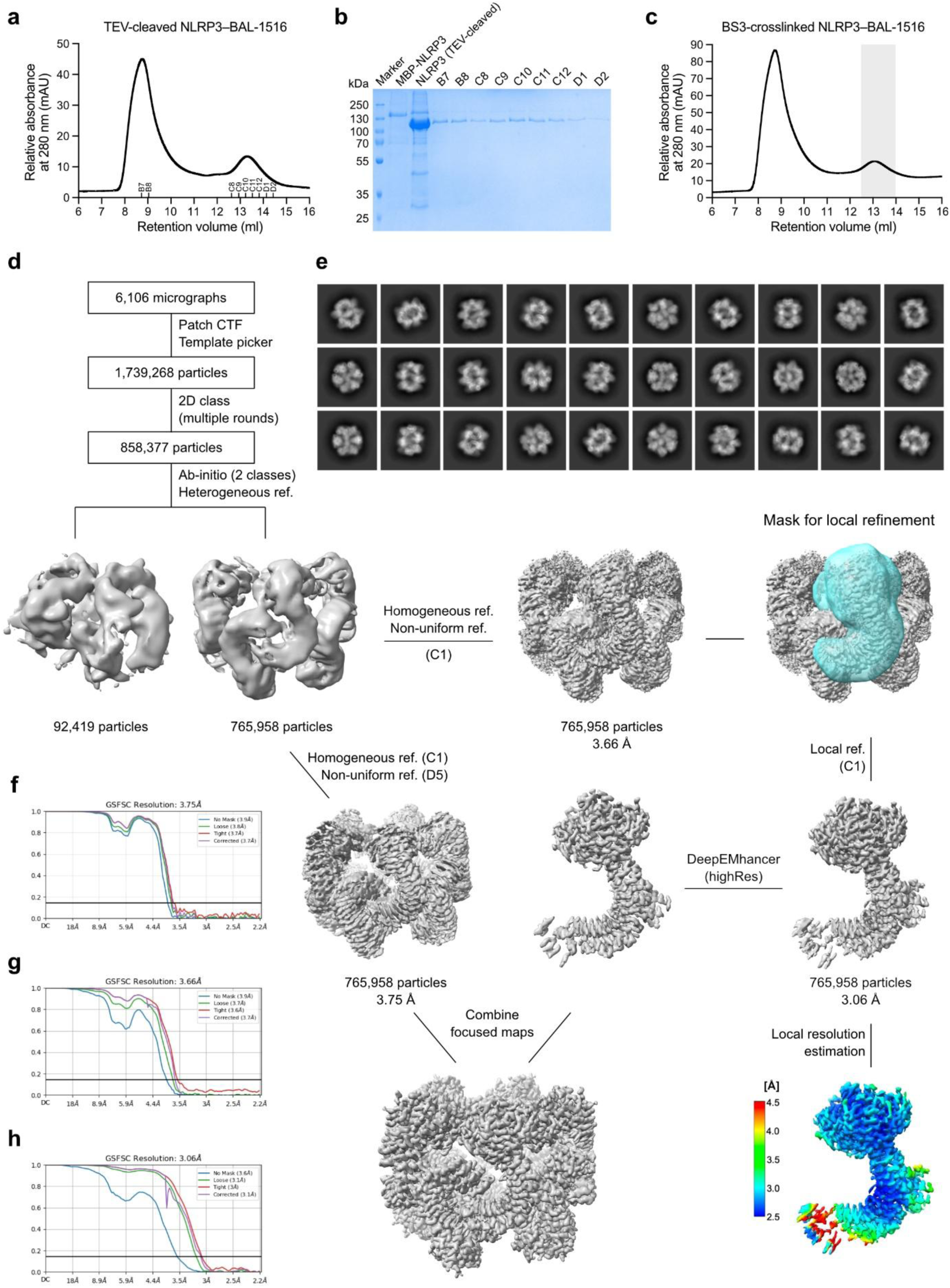
Cryo-EM data processing of the NLRP3–BAL-1516 complex. **a**, Size-exclusion chromatogram of NLRP3 with 10 µM BAL-1516 added during purification. The protein sample was TEV-digested prior to loading to a Superose 6 increase 10/300 GL column. **b**, Coomassie-stained SDS-PAGE analysis of NLRP3 size-exclusion fractions as indicated in panel a. **c**, Size exclusion chromatogram of NLRP3 with 10 µM BAL-1516 added during purification. The protein sample was TEV-digested and cross-linked with BS3 prior to loading to a Superose 6 increase 10/300 GL column. Fractions highlighted in grey were pooled and concentrated to 0.78 mg/ml for cryo-EM sample preparation. **d**, Cryo-EM reconstruction workflow in CryoSPARC v4.4.0 for human NLRP3 with BAL-1516. The particle picking and 2D classification rounds, followed by an ab-initio and heterogeneous refinement jobs with 2 classes yielded an initial volume representing the NLRP3 decamer based on 765,958 particles. Further global refinements with C1 symmetry yielded a 3.66 Å resolution map. Focusing mask was applied on one monomer for a local refinement, which improved the resolution to 3.06 Å. This local refinement map was further improved with DeepEMhancer (highRes). For the composite map of the decamer, D5 symmetry was imposed in the non-uniform global refinement job. This D5-symmetric map was used as a base map, onto which the locally refined monomers were projected in Phenix v1.21.2-5419-000. **e**, Final 2D classes used for the reconstruction. **f**, Gold-standard FSC curve for global refinement with D5 symmetry. **g**, Gold-standard FSC curve for global refinement with C1 symmetry. **h**, Gold-standard FSC curve for local refinement with the monomer mask. The monomer and decamer maps are deposited in the Electron Microscopy Data Bank under accession codes EMD-52874 and EMD-52908, respectively. The model coordinates are available at the Protein Data Bank under accession codes 9IHN and 9Q8V for the monomer and decamer structures, respectively.

**Extended Data Fig. 3.**
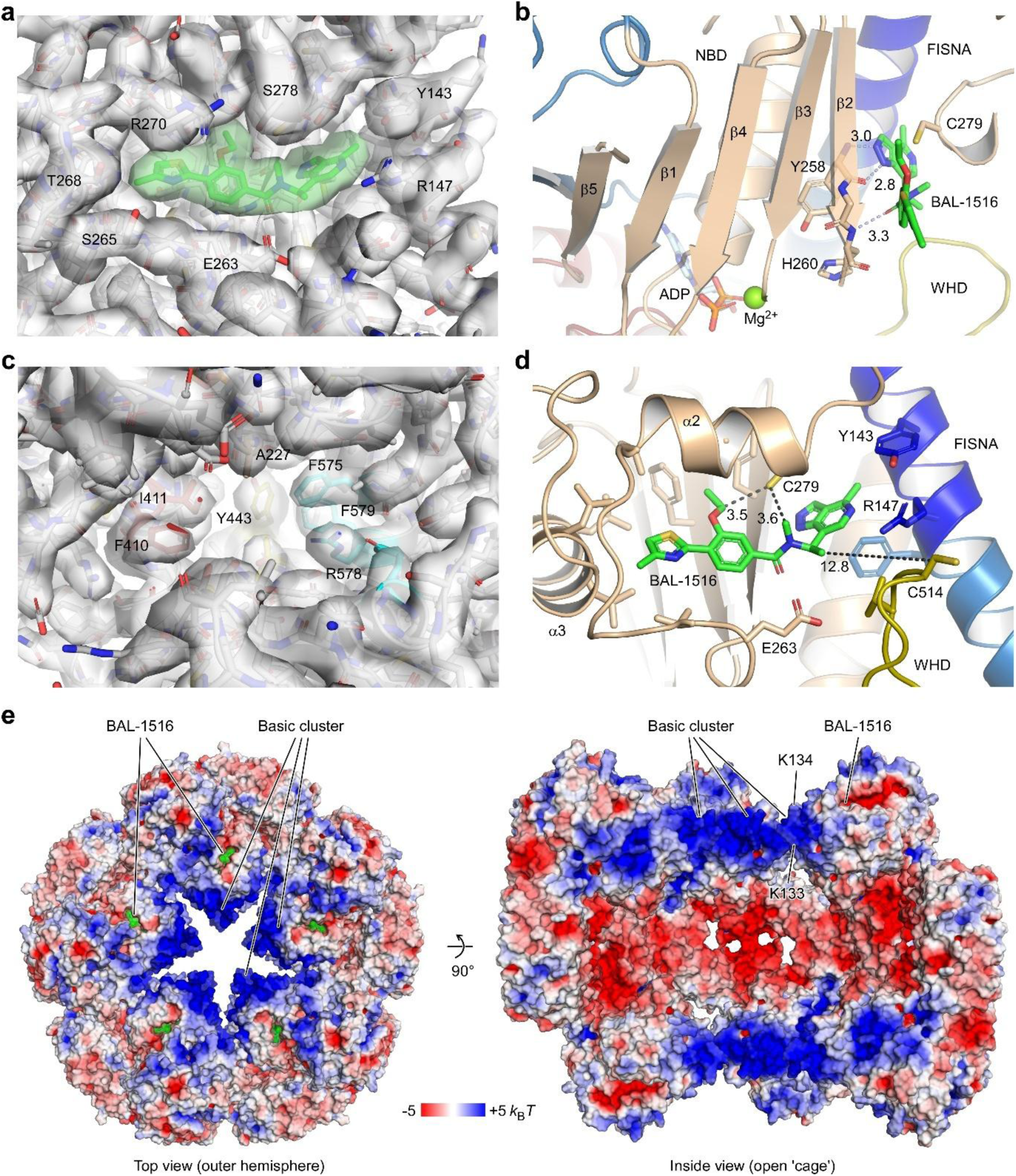
Details of the BAL-1516 interaction with NLRP3. **a**, Density map around BAL-1516 in the high-resolution monomer map displayed at 5 RMSD threshold. The density identifying BAL-1516 is coloured green. Residues of NLRP3 surrounding the compound are labelled. **b**, Residues Y258 and H260 at the tip of the terminal β-strand β2 of the NBD form three hydrogen bonds with the BAL compound in the manner of an antiparallel β-strand. This characteristic interaction with the β-sheet structure of the triple-ATPase domain is the defining feature of the indazole-based BAL compound binding mode. **c**, Display of the density map around the MCC950-binding site in the NLRP3–BAL-1516 complex structure reveals an empty site. **d**, Two cysteines of NLPR3 are in proximity to the BAL compound. C279 is in 3.5 or 3.6 Å distance to the ethoxy or methyl-aminocarbonyl groups of BAL-1516, respectively. C514 at the tip of the β-hairpin loop in the WHD is in 13 Å distance to the methyl group of the chiral centre. **e**, Display of the electrostatic surface potential shows the proximity of the BAL-1516 binding site to the basic cluster (131-147) of NLRP3. Whereas the basic cluster at the N-terminus of the FISNA forms a positively charged rim, the inner torus is negatively charged.

**Extended Data Fig. 4.**
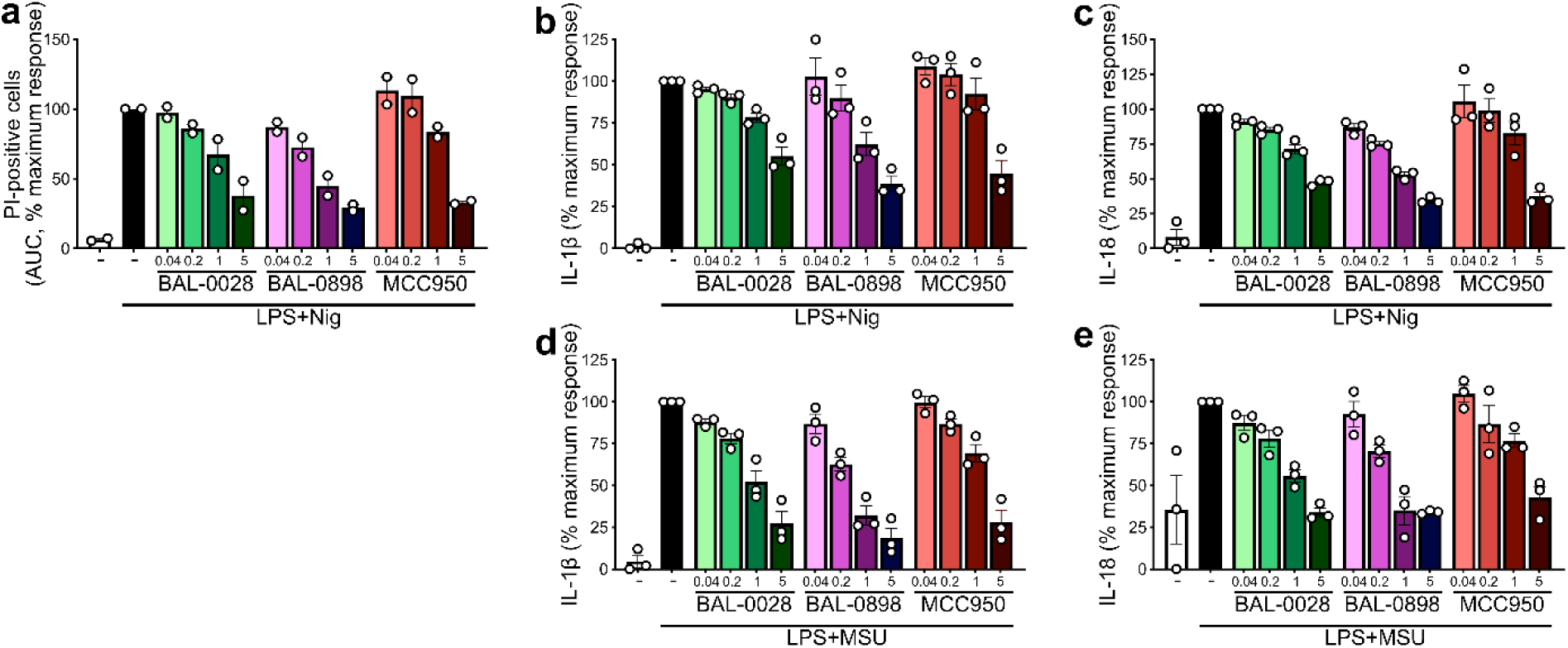
BAL compounds inhibit inflammasome activity in human primary monocytes and PBMCs. **a**, Dose-response measurements of the inhibition of pyroptosis by BAL-0028, BAL-0898 and MCC950 in human primary monocytes. Cells were primed with 40 ng/ml LPS for 4 h and stimulated with 5 µg/ml nigericin in the presence of the inhibitors at the indicated doses (in µM) for 2 h. **b-e**, Dose response measurements for BAL-0028, BAL-0898 and MCC950 inhibition of IL-1β (b, d) and IL-18 (c, e) release in human PBMCs. Cells were primed with 10 ng/ml LPS for 3 h and stimulated with 5 µg/ml nigericin for 2 h (b, c) or 100 µg/ml MSU for 3 h. Data plotted for mean ± s.e.m. of 2-3 independent experiments, each performed in 2-4 technical replicates.

**Extended Data Fig. 5.**
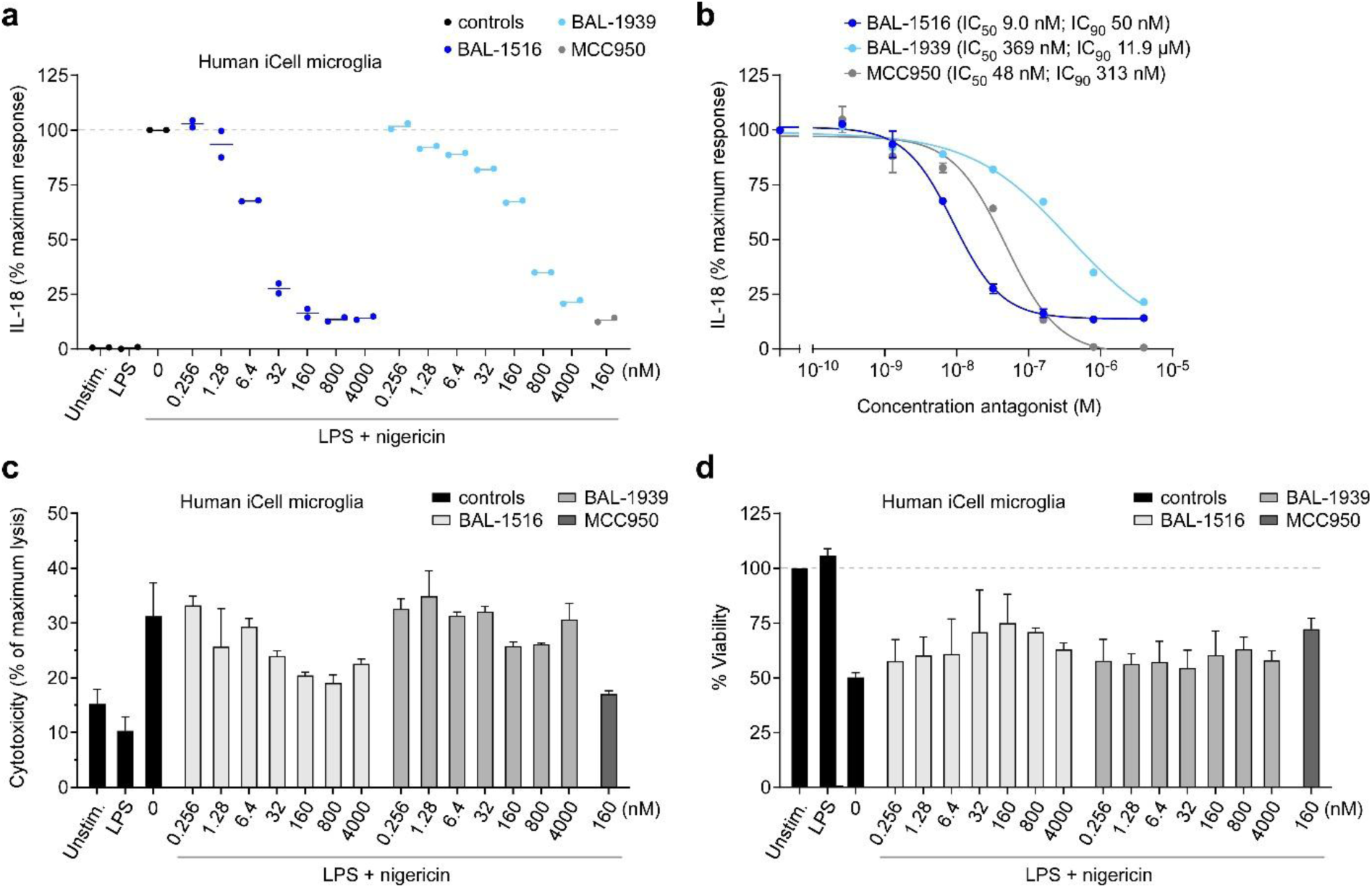
BAL-1516 is a potent inhibitor of NLRP3 inflammasome in human microglia. **a**, Inhibition of IL-18 release by BAL-1516 or the enantiomer BAL-1939 from iCell microglia primed with 200 ng/ml LPS for 4 h and stimulated with 10 µM nigericin for 30 min. **b**, Dose response curve for BAL-1516, BAL-1939 and MCC950 inhibition of IL-18 release in iCell Microglia. IC50s and IC90s were determined from mean maximum response compared with positive control. **c**, Effect of BAL-1516 and BAL-1939 on cytotoxicity in iPSC-derived microglia after LPS priming and nigericin stimulation as determined by LDH release. **d**, Cell viability of iPSC-derived microglia upon BAL-1516 or BAL-1939 treatment after LPS priming and nigericin stimulation as determined with the CellTiter-Blue assay.

**Extended Data Fig. 6.**
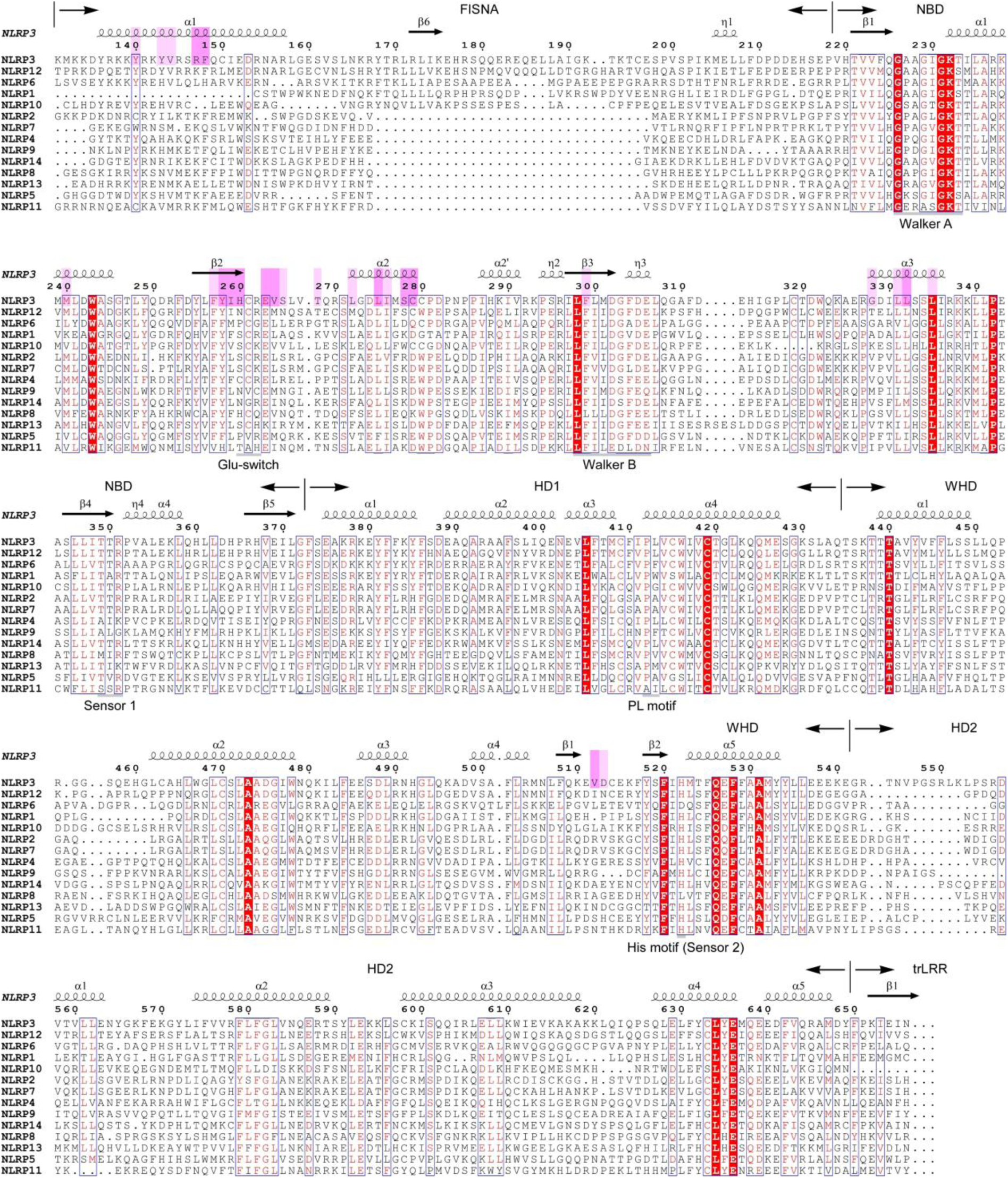
Mapping of the BAL-1516 binding site to human NLRPs. Residues in NLRP3 interacting with BAL-1516 are marked above the sequence. Light bars correspond to 1–10, dark bars to > 10 Å^2^ buried surface area, respectively. Secondary structure elements and subdomain boundaries are indicated at the top; common sequence motifs of STAND family triple-ATPases are marked at the bottom. Sequences of human NLRP proteins NLRP1 (UniProt accession number Q9C000), NLRP2 (Q9NX02), NLRP3 (Q96P20), NLRP4 (Q96MN2), NLRP5 (P59047), NLRP6 (P59044, Δ591-614), NLRP7 (Q8WX94), NLRP8 (Q86W28), NLRP9 (Q7RTR0), NLRP10 (Q86W26), NLRP11 (P59045), NLRP12 (P59046), NLRP13 (Q86W25) and NLRP14 (Q86W24) were aligned with Clustal Omega, and the secondary structure annotated with ESPript^36^. Sequences are ordered according to their degree of homology to NLRP3. The closest homolog of NLRP3, NLRP12, already contains changes of eight residues within the 26 residues that form the BAL compound binding site in NLRP3.

**Extended Data Fig. 7.**
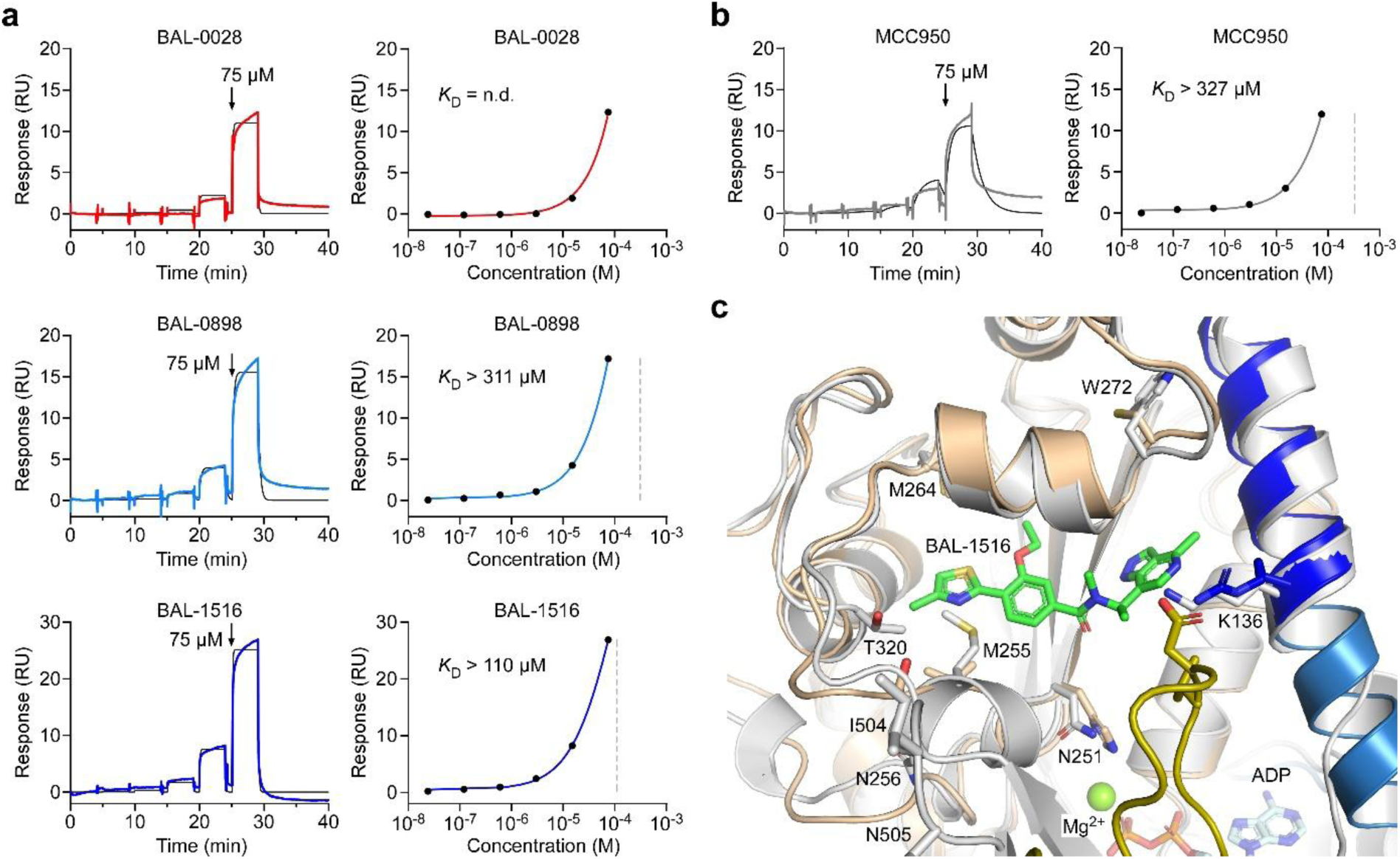
SPR analysis of the interaction of BAL compounds with NLRP12. **a**, SPR sensorgrams of human NLRP12 (122-679) with BAL compounds 0028, 0898 and 1516. Single-cycle kinetic experiments were performed with immobilized biotinylated NLRP12 as ligand and six injections of compounds as analytes with each injection increasing the concentration by a factor five to a maximal concentration of 75 µM (left panels). No dissociation constant (*K*D) could be determined (n.d.) for BAL-0028. BAL-0898 and BAL-1516 yielded *K*D’s larger than 100 µM as indicated by the dashed lines in the affinity plots (right panels). The interaction of BAL-0028, BAL-0898 and BAL-1516 with human NLRP12 is therefore at least 1,000-fold weaker than binding to human NLRP3. **b**, For comparison, the interaction of MCC950 to human NLRP12 was analysed using the same experimental setup. A dissociation constant larger than 300 µM was determined for NLRP12, which is in agreement with the high specificity of MCC950 for NLRP3. **c**, Superimposition of the NLRP3–BAL-1516 structure determined here (9IHN, coloured) with the AlphaFold2 model of human NLRP12 (grey). Residues in NLRP12 which differ from residues in the BAL compound binding site in NLRP3 are shown as sticks and labelled by the amino acid. Particularly the larger residues M255 in NLRP12 (V264 in NLRP3), T320 in NLRP12 (G328 in NLRP3), and W272 in NLRP12 (C280 in NLRP3) restrict the space of the compound binding site, explaining its low potency for NLRP12.

**Extended Data Table 1.** Cryo-EM data collection, refinement and validation statistics.

|  |  |  |
| --- | --- | --- |
| Map and model | NLRP3 monomer<br>bound to BAL-1516 | Composite map of the NLRP3<br>decamer bound to BAL-1516 |
| EM density map accession code | EMDB-52874 | EMDB-52908 |
| PDB accession code | 9IHN | 9Q8V |
| **Data collection and processing** |  |  |
| Magnification | 105,000 | 105,000 |
| Voltage (kV) | 300 | 300 |
| Electron exposure (e-/Å<sup>2</sup>) | 51.7 | 51.7 |
| Defocus range (µm) | -2.2 to -0.8 | -2.2 to -0.8 |
| Pixel size (Å) | 0.82 | 0.82 |
| Symmetry imposed | C1 | D5 |
| Initial particle images (no.) | 1,739,268 | 1,739,268 |
| Final particle images (no.) | 765,958 | 765,958 |
| Map resolution (Å) | 3.06 | 3.06 |
| FSC threshold 0.143 |  |  |
| Map resolution range (Å) | 2.41 to 37.87 | 2.41 to 37.87 |
| **Refinement** |  |  |
| Initial model used (PDB code) | 7PZC | 7PZC |
| Map sharpening *B* factor (Å<sup>2</sup>) | Map sharpened with<br>DeepEMhancer (highRes) | Map sharpened with<br>DeepEMhancer (highRes) |
| Model composition |  |  |
| Non-hydrogen atoms | 6,713 | 67,130 |
| Protein residues | 833 | 8,330 |
| Ligands | ADP, Mg<sup>2+</sup><br>BAL-1516 | 10 ADP, 10 Mg<sup>2+</sup><br>10 BAL-1516 |
| *B* factors (Å<sup>2</sup>) |  |  |
| Protein | 57.88 | 59.49 |
| Ligand | 15.65 | 16.64 |
| R.m.s. deviations |  |  |
| Bond lengths (Å) | 0.003 | 0.002 |
| Bond angles (°) | 0.601 | 0.528 |
| Validation |  |  |
| MolProbity score | 1.56 | 1.40 |
| Clashscore | 3.35 | 4.63 |
| Poor rotamers (%) | 2.25 | 1.47 |
| Ramachandran plot |  |  |
| Favored (%) | 97.09 | 97.84 |
| Allowed (%) | 2.79 | 2.04 |
| Disallowed (%) | 0.12 | 0.12 |

